# Lipid nanodiscs as a template for high-resolution cryo-EM structures of peripheral membrane proteins

**DOI:** 10.1101/2023.03.07.531120

**Authors:** Kevin S. Cannon, Reta D. Sarsam, Tanita Tedamrongwanish, Kevin Zhang, Richard W. Baker

## Abstract

Peripheral membrane proteins are ubiquitous throughout cell biology and are required for a variety of cellular processes such as signal transduction, membrane trafficking, and autophagy. Transient binding to the membrane has a profound impact on protein function, serving to induce conformational changes and alter biochemical and biophysical parameters by increasing the local concentration of factors and restricting diffusion to two dimensions. Despite the centrality of the membrane in serving as a template for cell biology, there are few reported high-resolution structures of peripheral membrane proteins bound to the membrane. We analyzed the utility of lipid nanodiscs to serve as a template for cryo-EM analysis of peripheral membrane proteins. We tested a variety of nanodiscs and we report a 3.3 Å structure of the AP2 clathrin adaptor complex bound to a 17-nm nanodisc, with sufficient resolution to visualize a bound lipid head group. Our data demonstrate that lipid nanodiscs are amenable to high-resolution structure determination of peripheral membrane proteins and provide a framework for extending this analysis to other systems.

## Introduction

The endomembrane network is a defining feature of eukaryotic cell biology. Virtually every cellular process requires some form of membrane engagement or membrane remodeling, including division, migration, and vesicle trafficking. While the physicochemical properties of membranes — such as 2D planarity, curvature, fluidity, and charge — undoubtedly define and control cellular reactions^1^, the membrane is often omitted from structural and biochemical studies because of the technical challenges it poses. This severely limits our understanding of the numerous biological processes that are dependent on membranes to drive protein complex assembly, induce conformational changes, and serve as a platform for downstream signaling cascades.

While there are thousands of peripheral membrane proteins in eukaryotes^2^, only a modest number of structures of proteins bound to the surface of the membrane have been reported. Membrane-containing samples are generally refractory to X-ray crystallography, and only with recent advances in cryo-electron microscopy (cryo-EM) has structure determination of more heterogeneous samples been feasible. This includes an explosion in the number of reported structures of integral membrane proteins embedded in lipid micelles, nanodiscs, or other amphipathic polymers that stabilize lipid-embedded proteins^3^. In contrast, there are relatively few structures of peripheral membrane proteins bound to membranes. Examples of high-resolution structures (>4 Å) are restricted to filamentous or helical polymers that tubulate liposomes and enforce a rigid architecture on the membrane that aids in structure determination. Additionally, there are a number of tomographic reconstructions of peripheral membrane assemblies, notably several vesicle coats including COPI^4^, retromer^5^, and clathrin^6^, although these are also oligomeric assemblies and are generally limited to moderate resolutions (6 - 10 Å). However, most peripheral membrane proteins do not form filaments or large oligomeric assemblies and currently there are no high-resolution structures of a non-symmetric protein bound to a membrane.

We sought to test the utility of lipid nanodiscs^7^, a common tool for studying the structure and function of integral membrane proteins, as scaffolds for high-resolution structural characterization of peripheral membrane proteins. Nanodiscs are disc-shaped lipid bilayers enclosed by two copies of an amphipathic membrane scaffold protein (MSP), derived from the lipid-binding protein Apolipoprotein A-1^8^. Variations in the length of MSP sequences have enabled experimentalists to readily tune the diameter, and thus, available membrane area, of nanodiscs with nanometer-scale precision. To test the feasibility of this approach, we chose to study the clathrin coat component AP2^9^. As a constitutive part of the endocytic clathrin coat, AP2 serves as the primary regulatory hub of endocytosis and functions during the earliest stages of vesicle formation^9,10^ (Supp. Fig. 1). AP2 function is largely dictated by a series of large-scale conformational changes, which expose binding sites for phospholipids, two unique classes of trans-membrane cargo, and clathrin^6,11–13^. However, this model is largely based on X-ray and cryo-EM structures of soluble components^6,11–17^, limited proteolysis experiments^18,19^, and a moderate resolution structure of the fully assembled AP2-clathrin coat on liposomes^6^. Additionally, endocytic regulators have also been proposed to induce membrane-bound conformations of AP2^16^ that are distinct from the conformation seen in vesicles^6,11^. High-resolution nanodisc-AP2 structures could therefore be essential for deciphering the full conformational cycle of coat assembly as AP2 engages membranes, interacts with regulators, binds trans-membrane cargo, and finally templates assembly of the clathrin cage.

Here, we report several single particle cryo-EM structures of AP2 bound to lipid nanodiscs, including a 3.3 Å cryo-EM structure of AP2 bound to a lipid nanodisc containing the plasma membrane-enriched lipid phosphatidylinositol-4,5-bisphosphate (PIP_2_) and a lipidated peptide bearing the YxxΦ endocytic motif of Tgn38. We find that nanodisc diameter strongly influences the conformation and overall resolution of the final reconstructions. Small nanodiscs (∼9-13 nm) lead to moderate resolution reconstructions or non-native conformations, whereas larger nanodiscs (∼17 nm), lead to high-resolution reconstructions in the native conformation. Consistent with the first reported crystal structure of AP2 bound to a YxxΦ cargo peptide^11^, and a more recent tomographic reconstruction of AP2 in assembled clathrin coats^6^, we find AP2 in the canonical open conformation when bound to a cargo-containing lipid nanodiscs. We also observe different conformations of AP2 when comparing nanodiscs with and without cargo peptides, indicating that cargo binding causes a conformational rearrangement in AP2. The quality of our highest-resolution model is sufficient to visualize a PIP_2_ headgroup engaging a known lipid-binding site on the *α* subunit. Our data therefore provide new insights into the earliest stages of vesicle assembly by describing in molecular detail the initial binding of AP2 to PIP_2_ membranes, followed by a subsequent conformational change upon engagement with trans-membrane cargo. Our approach to determining cryo-EM structures of peripheral membrane proteins should be broadly applicable to a range of biological systems, elucidating previously overlooked aspects of cell biology that take place at membrane surfaces.

## Results

### AP2-nanodisc complex reconstitution and purification

AP2 contains three PIP_2_-binding sites that all contribute meaningfully to membrane recruitment^11^. We reasoned that the size of the nanodisc surface would be critical to ensure that all membrane binding sites could be accommodated without restriction (Fig. 1A-B). Based on published reports, we purified and assembled nanodiscs of varying predicted diameters using the following MSP variants: MSP1 (9 nm), MSP1E3D1 (13 nm), and MSP2N2 (17 nm). All nanodiscs were assembled with a lipid mixture containing 75 mol% DOPC, 15 mol% DOPS, 10% PIP_2_. Where indicated, lipidated YxxΦ-cargo with a FITC moiety was included at 2 mol%, and DOPC was reduced to 73 mol%. We used mass photometry^20^ to characterize the molecular weight for each nanodisc, and observed a mass distribution indicative of the predicted molecular weight (Supp. Fig. 2A,B). We used negative stain transmission electron microscopy (TEM) to verify the diameter of each nanodisc, which was consistent with previous reports^21,22^ (Fig. 1C, Supp. Fig. 2C,D).

**Figure 1.**
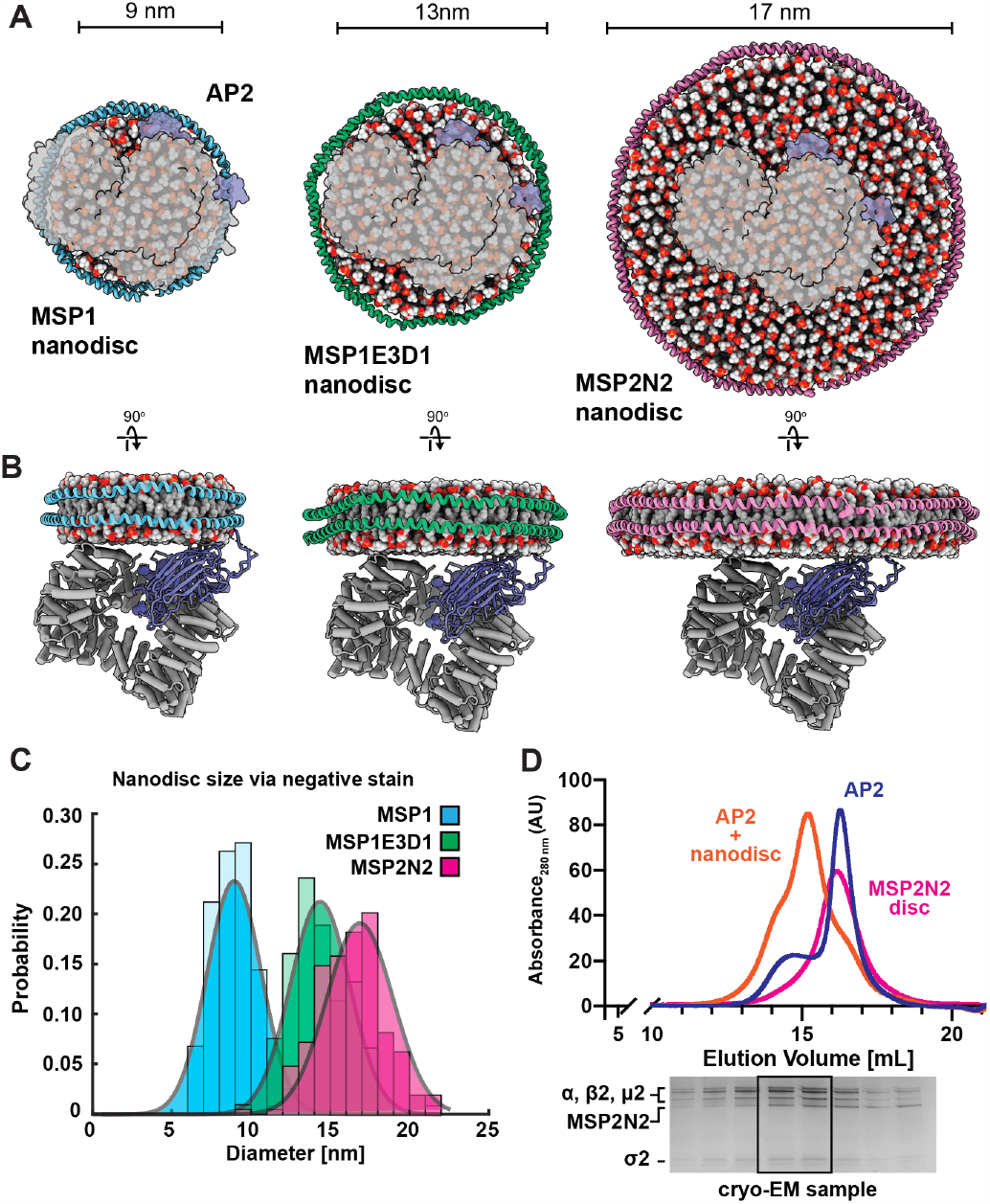
Reconstitution of AP2-nanodisc complexes. (A) Schematic highlighting a top-down view of AP2 (grey and blue) bound to nanodiscs of various sizes. AP2 and nanodiscs are drawn to scale. Nanodisc PDB models are idealized representations made using the CHARMM-GUI. (B) Side-on view of predicted AP2-nanodisc complexes, shown in cartoon representation. (C) Nanodisc diameter distribution obtained from negative stain TEM. Gaussian fitting was used to calculate the mean. N > 100 nanodiscs per condition. (D) Representative gel filtration profiles of AP2 (blue trace), MSP2N2 nanodiscs (pink trace), and AP2-nanodisc complexes (orange trace). The gel corresponds to the elution profile of the AP2-nanodisc complex (orange trace) highlighting the co-elution of AP2 with the MSP2N2 nanodisc.

AP2 has intrinsic, albeit moderate, affinity for PIP_2_-containing membranes, which is strongly enhanced by the presence of trans-membrane cargo^11,23^. Surface plasmon resonance (SPR) analysis shows an apparent affinity of 7 μM for PIP_2_ liposomes alone compared to 320 nM for PIP_2_ + YxxΦ cargo^11^, an increase of over an order of magnitude. With this in mind, we first attempted to form AP2-nanodisc complexes using nanodiscs containing PIP_2_ and a lipidated YxxΦ cargo peptide from Tgn38, a nearly identical construct used in other studies^6,11^. We performed gel filtration binding assays between AP2 and PIP_2_-cargo nanodiscs of three diameters (9, 13, 17 nm). We observed strong binding for all three nanodisc diameters tested, as shown by a large shift in the elution profile and visualized by SDS-PAGE (Fig. 1D; Supp. Fig. 3, right).

While AP2 binds to PIP_2_ membranes absent other factors, these experiments are typically performed using large membrane surfaces such as liposomes tethered to an SPR sensor or liposome flotation assays. Considering that binding of AP2 to membranes is likely to be strongly enhanced by the number of available binding sites on a liposome, we were uncertain if AP2 would bind to nanodiscs in a stoichiometric manner require d for structural analysis. For example, a 100 nm liposome has nearly 150 times the available surface area of our largest 17 nm nanodisc. Considering an apparent affinity of 7 μM for PIP_2_ liposomes, we predicted sub-stoichiometric binding of AP2 to PIP_2_ nanodiscs. Surprisingly, we observe robust, stoichiometric binding for PIP_2_ nanodiscs, as observed via gel filtration chromatography (Supp. Fig. 3, left). These results are indistinguishable from our analysis with nanodiscs containing a YxxΦ cargo motif (Supp. Fig. 3, right). This result suggests that proteins with low or moderate affinities for membranes should be empirically tested for binding to nanodiscs.

### High resolution AP2 structures using lipid nanodiscs

AP2 has previously been shown to be in a compact, inactive conformation in its soluble state^14^, and in an open, active conformation when bound to cargo peptides^11,16,24^ or cargo-containing membranes^6^. The main structural change is a large displacement of the μ2 C-terminal domain (μ2-CTD) (Supp. Fig 1), which contains both a PIP_2_-binding site^14^ and the YxxΦ-cargo binding site^25^. To see if nanodiscs could recapitulate known membrane-induced allostery, we took our various AP2-nanodisc complexes and collected single particle cryo-EM data on a 200 keV Talos Arctica cryo-TEM, yielding six total datasets: 3 nanodiscs of varying diameter, either “empty” where the only contact with AP2 is binding to PIP_2_ headgroups or “with-cargo”, where a lipid-conjugated YxxΦ-cargo peptide that binds to the μ2-CTD of AP2 is incorporated into the nanodisc (summarized in Table 1). For all datasets, 2D classification shows an “open” conformation of AP2, where the μ2-CTD is ejected from the closed conformation, demonstrating that the membrane is sufficient to allosterically activate AP2 (Supp. Fig. 4-6). Extra density corresponding to the nanodisc at known membrane binding sites is also visible (Supp. Fig. 4-6). Following extensive 2D and 3D classification, we obtained final reconstructions of nanodisc-bound AP2 complexes with resolutions ranging from 3.3 to 6.8 Å. To improve the resolution of our best reconstruction, three local refinements for the MSP2N2 + YxxΦ-cargo condition were performed (Supp. Fig. 7).

**Table 1.**
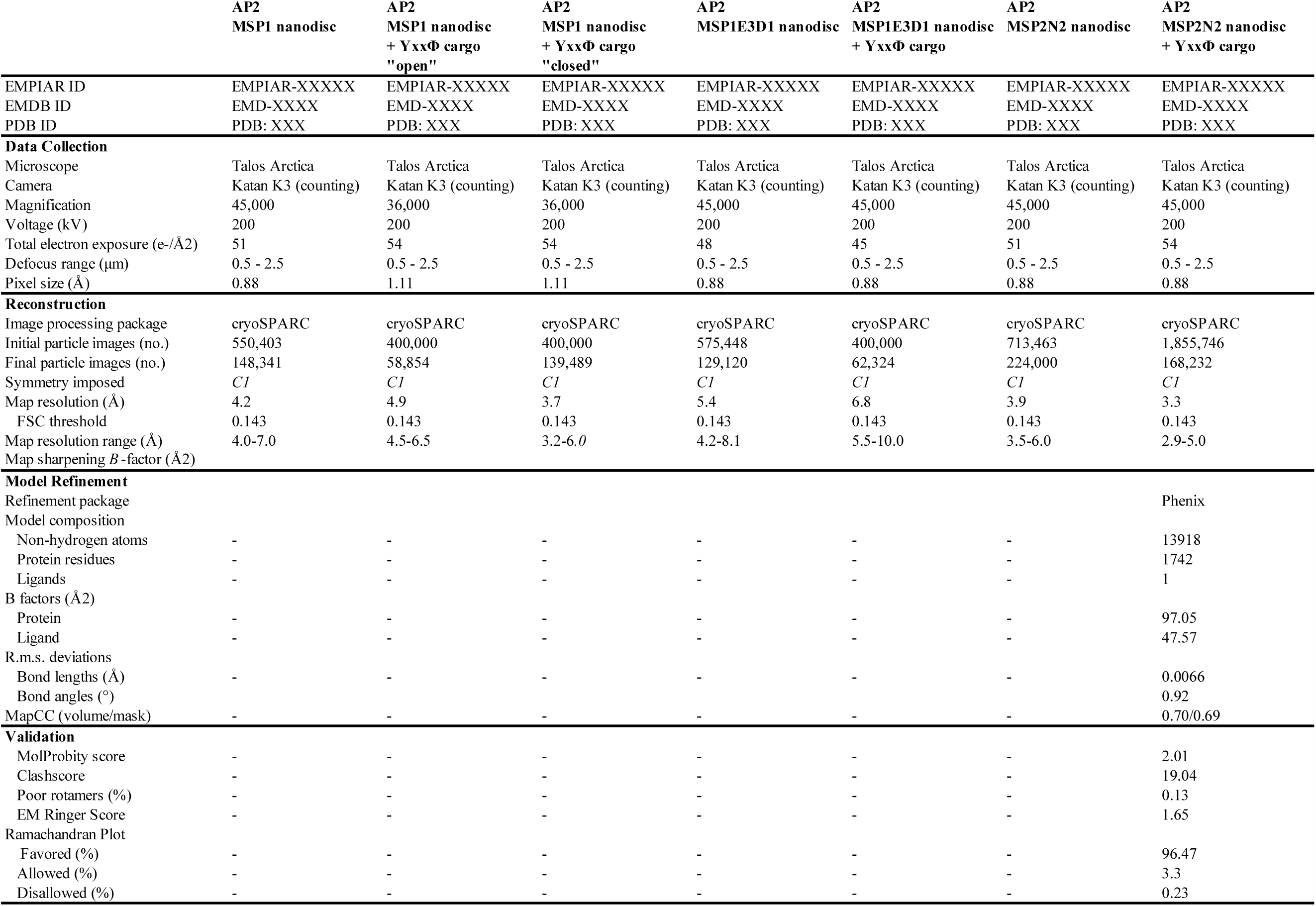
Cryo-EM data collection, refinement, and validation statistics.

### Overall resolution and conformation is dependent on nanodisc diameter

Although we observe an open AP2 conformation using all na nodisc sizes tested, their overall resolution and conformation is dependent on both nanodisc diameter and the presence of the YxxΦ-cargo peptide (Fig. 2 and Supp. Fig. 4-6). Our data show that on nanodiscs lacking cargo, we observe density for *α*/β2/σ2 subunits and the μ2 N-terminal domain (NTD), which has canonically been referred to as the AP2 “bowl” (Fig. 2A). Importantly, we do not observe density for the μ2-CTD, suggesting that this domain does not adopt a fixed location in the absence of cargo peptide. As the μ2-CTD contains both a PIP_2_ and cargo binding site, it is unclear whether this domain is free in solution, or is still bound to the nanodisc, albeit without a fixed location relative to the bowl of AP2. This is in contrast to AP2 structures on nanodiscs that contain cargo peptide, as we observe density for the μ2-CTD on two of three nanodisc sizes tested (Fig. 2B). These data suggest that that the conformational landscape sampled by the μ2-CTD of AP2 can be tuned by the binding YxxΦ-cargo motifs.

**Figure 2.**
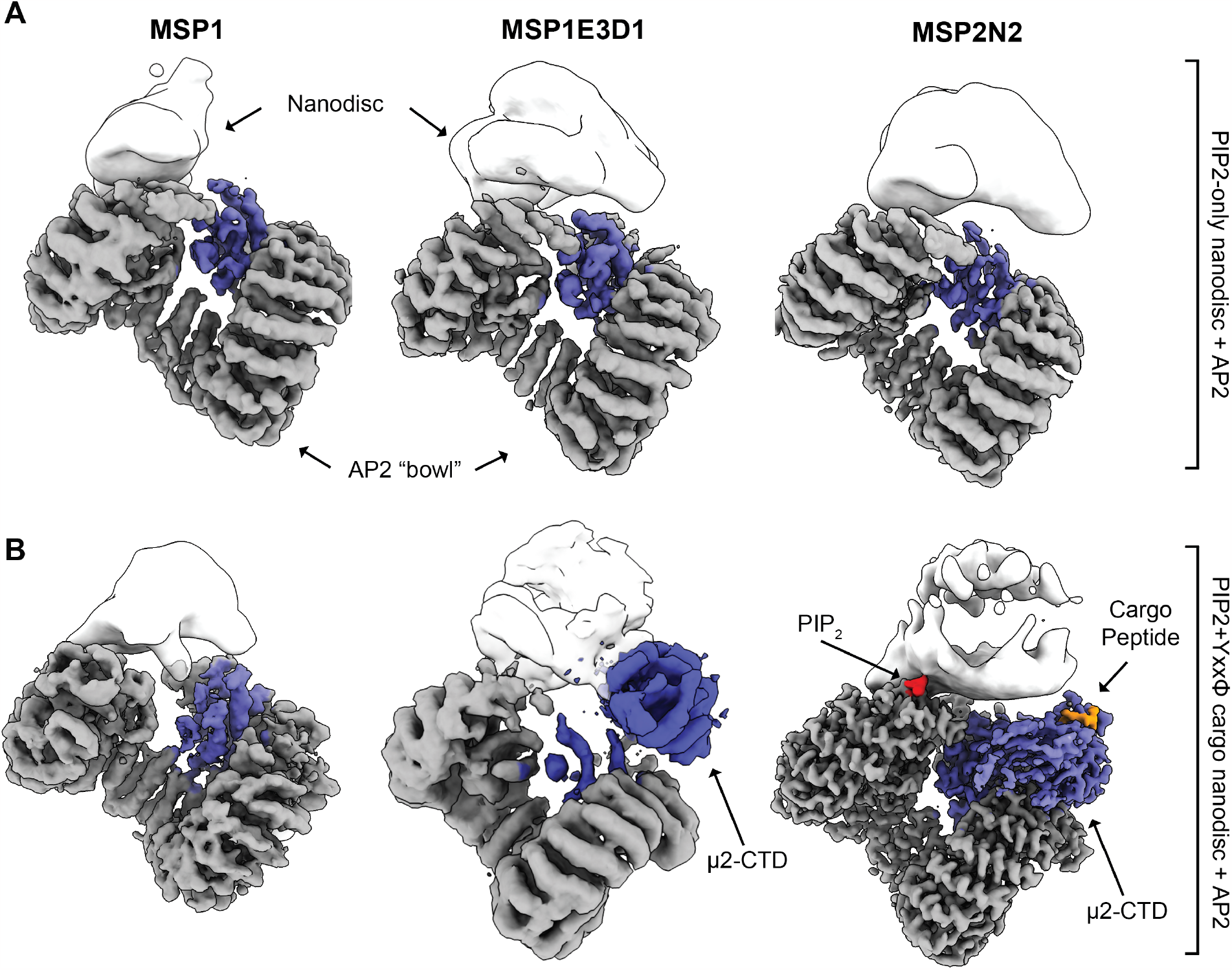
Cryo-EM structures of AP2-nanodisc complexes. (A) Cryo-EM structures of AP2 bound to the three nanodisc constructs are shown. Maps were manually segmented and the nanodisc density was gaussian filtered in UCSF Chimera. Nanodisc density is shown in white, and AP2 is colored grey (*α*/β2/σ2) and blue (μ2). (B) Cryo-EM structures of three nanodisc constructs containing a lipidated YxxΦ-cargo peptide are shown. Segmentation, filtering, and coloring were performed as in (A). All maps shown were sharpened using deepEMhancer^55^.

The degree to which the μ2-CTD could be resolved and its position relative to the membrane was dependent on nanodisc size (Supp. Fig. 8). The largest nanodiscs (MSP2N2; ∼17 nm) gave the highest overall resolution as well as the best resolved μ2-CTD of AP2 (Fig. 2). In this reconstruction, the bound YxxΦ-cargo motif is also visible. In contrast, we did not observe any particles with a fixed μ2-CTD when using the smallest nanodisc (MSP1; ∼9 nm). This is perhaps due to insufficient membrane surface to accommodate both *α* and μ2. Furthermore, we observe that the 13 nm MSPE3D1 + YxxΦ-cargo nanodisc sample is our lowest resolution structure and that AP2 is in a non-native conformation (Supp. Fig. 8). In this structure, the μ2-CTD is brought closer to the PIP_2_ binding site on *α*, essentially “pinching” AP2 relative to the canonical conformation. As AP2 binds to the membrane through placement of five membrane binding sites (three PIP_2_ sites and two for trans-membrane cargo) that span ∼40-50 Å from one another in a coplanar geometry, we speculate that differences in achievable resolution are due to how well the available membrane area accommodates these binding sites (Supp. Fig. 8). This consideration for resolving high-resolution structures likely applies to any peripheral membrane complex with multiple membrane binding sites.

### PIP_2_ headgroups are resolvable within AP2 binding pockets on native membranes

The molecular basis for AP2 binding to PIP_2_ is largely based on the first X-ray crystal structure of the full AP2 complex, which was co-crystallized with the soluble PIP_2_ analog, inositol hexaphosphate (IP6)^14^. In this structure, a bound IP6 molecule is observed at a crystal contact between the *α* and μ2 subunits of symmetry - related AP2 molecules. Both the *α* and μ2 binding sites rely on a series of basic residues that extend “fingers” into the solvent to coordinate the negative charge of the IP6 molecule. In our highest resolution structure, we observe a bound PIP_2_ head group in the *α* binding site (Fig. 3A). In our structure, the inositol ring binds in a perpendicular orientation relative to the surface of *α*, with a pair of Lysine and Tyrosine “fingers” coordinating each phosphate. In contrast, the inositol ring of IP6 in the original AP2 crystal structure shows a planar geometry, with three of the six phosphate groups being coordinated by residues in *α* (Fig. 3B). Several residues in the IP6 structure that contact the phosphate groups do not show the same interaction in our PIP_2_ structure (compare Fig 3A and B). Importantly, AP2 has also been reported to bind another phospholipid, phosphatidylinositol (3,4,5)-triphosphate (PIP_3_)^26,27^. We propose that in the case of AP2, the use of IP6 might best mimic the PIP_3_-bound state, as it engages more residues in AP2 than PIP_2_. The biological importance of our observed conformational changes is unclear. Overall, our observation of a clearly resolved lipid headgroup bound to AP2 in the presence of a native membrane demonstrates that nanodiscs can be used as a platform to probe the molecular and structural details of protein-lipid interactions.

**Figure 3.**
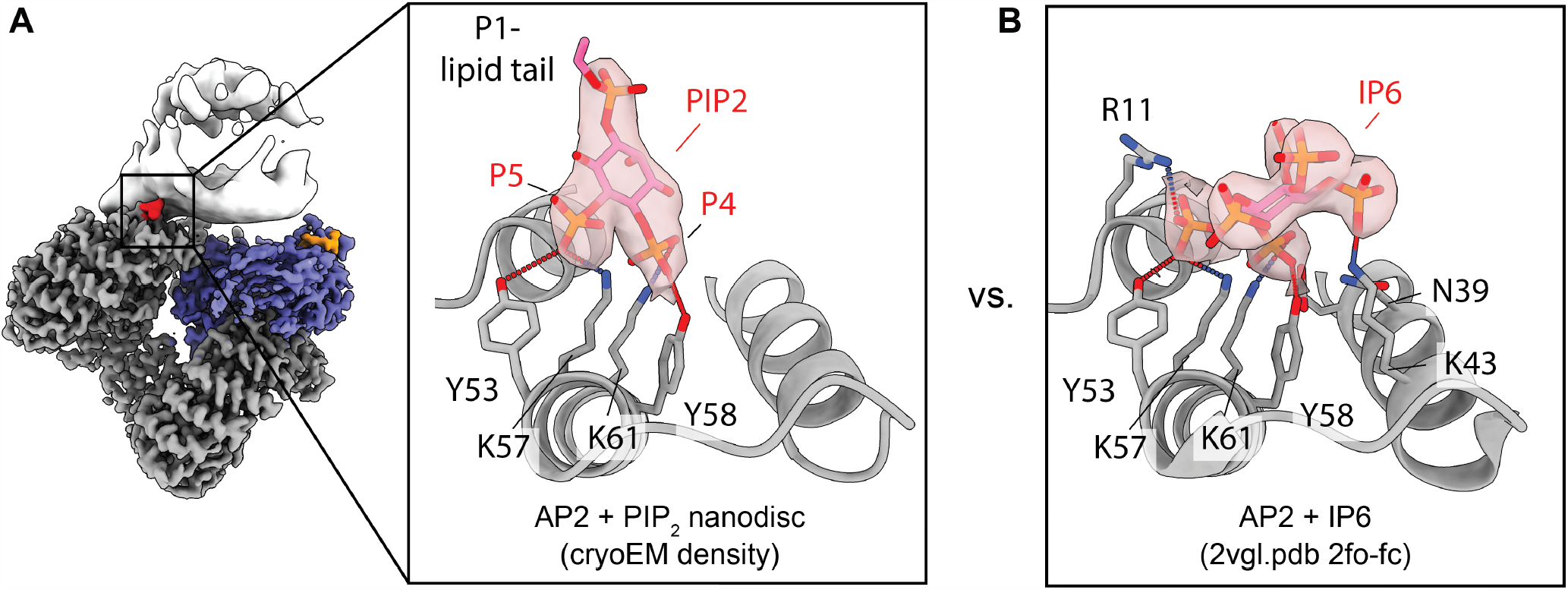
Comparison of PIP_2_-binding pocket on *α*. **(A)** The PIP_2_-binding pocket of *α* is shown for the MSP2N2 nanodisc structure. Density corresponding to PIP_2_ is shown with a transparent surface. Residues that directly contact the ligand are show in atom representation. (B) The PIP_2_-binding pocket of AP2 + IP6 (2VGL.pdb^14^, right) is shown. Density corresponding to IP6 is shown with a transparent surface. Residues that directly contact the ligand are show in atom representation.

## Discussion

Our data represent the first high-resolution structure of a peripheral membrane protein bound to a lipid nanodisc. To the best of our knowledge, the only other example of a similar complex is a moderate resolution structure of the clotting factor prothrombinase on a phosphatidylserine nanodisc^28^. More broadly, we lack structures of proteins bound to the surface of a membrane, limiting our understanding of how this ubiquitous biological surface affects protein function. The two most common classes of membrane-bound structures of peripheral membrane proteins are reconstructions of proteins decorating lipid nanotubes or GUVs and NMR ensembles of proteins bound to lipid nanodiscs. For example, a variety of vesicle coats^4,6,29^, viral capsid proteins^30,31^, and ESCRT polymers^32,33^ have been imaged on nanotubes/liposomes/GUVs, as well as the endocytic regulators Sla2-Ent1^34^ and the blood coagulation protein Factor VIII^35^. Other examples include BAR-domain containing proteins^36,37^ and the dynamin-like protein EHD4^38^. In general, these all form oligomeric assemblies that remodel or enforce a preferred geometry on the membrane. The few examples of high-resolution protein-membrane structures are almost exclusively polymers or filaments that can be analyzed using helical reconstruction methods, like dynamin^39^ and ESCRT^32,33^. A smaller number of membrane-bound structures that do not tubulate have been described, like 2D crystals of focal adhesion kinase (FAK) on lipid monolayers^40^ and a pore-forming complex, Macrophage expressed gene 1 (MPEG-1), in the pre-pore form bound to the surface of liposomes^41^. In contrast, NMR analysis of peripheral membrane proteins using nanodiscs usually focuses on small (<30 kDa) proteins like the GTPase KRAS^42–44^ and single alpha helices that insert into the lipid bilayer^45^.

Nanodiscs were originally engineered to stabilize biologically active conformations of otherwise insoluble integral membrane proteins. By rendering integral membrane proteins soluble within a native membrane environment, nanodiscs have been paramount for elucidating biophysical and structural mechanisms of vital cellular processes like G-protein coupled receptor signaling cascades^46^. Our results expand the nanodisc toolkit by determining high-resolution cryo-EM structures of the peripheral membrane protein AP2 using traditional single-particle techniques. While the presence of the nanodisc essentially acts as a disordered domain that increases the noise of each individual particle, we observe local resolutions up to 3 Å, including features like a bound PIP_2_ lipid headgroup. Our methodology therefore provides a framework to bridge the gap between high-resolution structures of soluble proteins via X-ray crystallography or single-particle cryo-EM, and moderate resolution structures of membrane-bound complexes using cryo-ET (Fig. 4).

**Figure 4.**
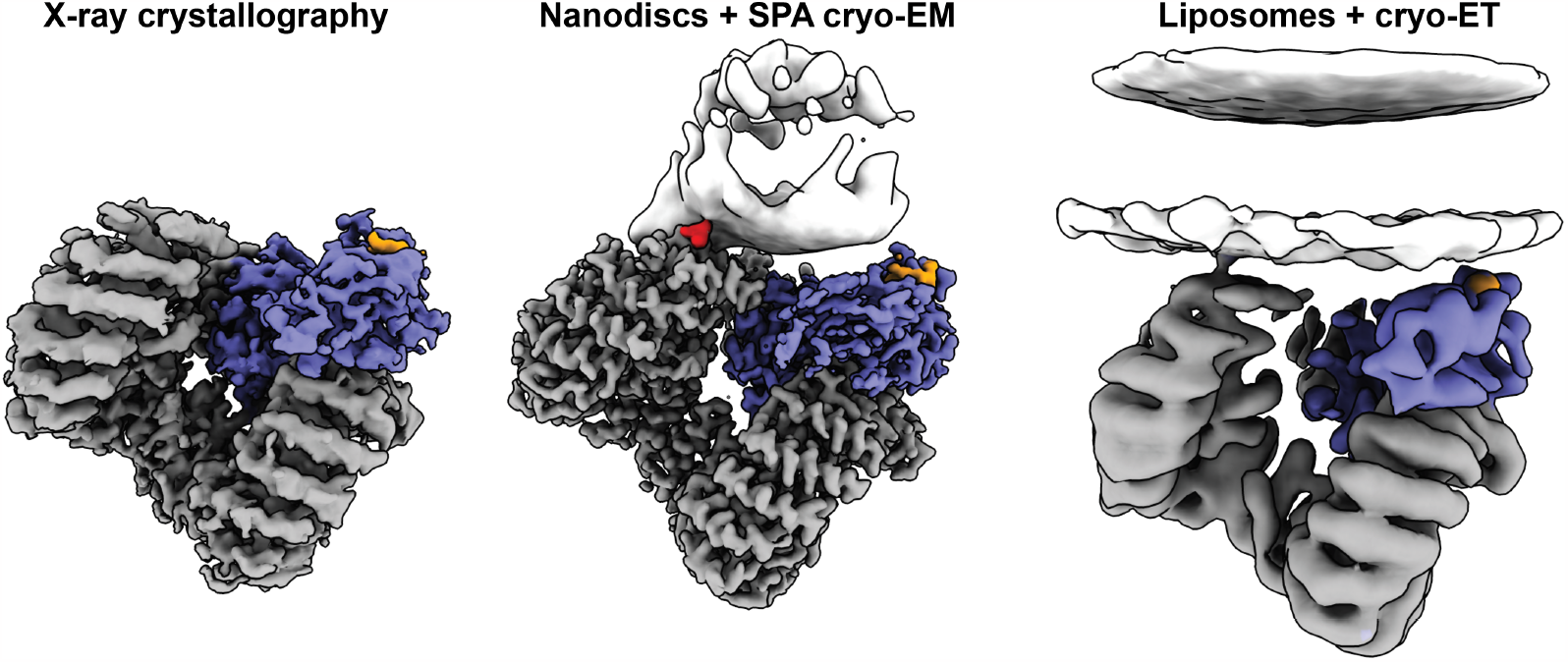
Comparison of three AP2 + YxxΦ-cargo structures. Three structures of AP2 bound to the YxxΦ-cargo motif of Tgn38 are shown. Left, the X-ray crystal structure of AP2 and a soluble cargo peptide (2XA7.pdb^11^). Middle, the cryo-EM structure of AP2 on a PIP_2_-containing MSP2N2 nanodisc with lipidated cargo peptide (this study). Right, the cryo-ET structure of AP2 on PIP_2_-containing liposomes with lipidated cargo peptide (EMD-10748^6^).

Our data suggest that nanodisc diameter and by extension the available membrane surface area is an important consideration for the success of structure enough affinity to observe stoichiometric binding (Supp. Fig. 3, left) at concentrations below the apparent affinity reported by other groups^11^. When trying to form stoichiometric complexes for analysis by cryo-EM, affinities with K_D_s in the low μM range are especially problematic, as the concentrations used to make grids are often in this concentration regime, which would determination. Based on our observation that 9-nm nanodiscs (MSP1) were too small to accommodate all membrane binding domains of AP2 (Supp. Fig. 8C, left), and that 13-nm nanodiscs (MSP1E3D1) could accommodate all membrane binding domains but that AP2 was “pinched” into a non-native conformation (Supp. Fig. 8C, middle), it seems paramount that the selected nanodisc have sufficient membrane area such that the all membrane-binding sites can be satisfied in a native conformation. Importantly, based on the predicted diameter of a 13-nm nanodisc, there should theoretically be sufficient space for AP2 to bind natively, but it did not lead to a high-resolution reconstruction. As it has been shown^47^ that the center of nanodiscs are thicker than the edges, and that the membrane properties like fluidity and disorder are also non-uniform, it seems likely that a significant excess of available membrane space might be ideal to accommodate peripheral membrane proteins for structural determination. Based on our results, a likely starting guideline is to use a nanodisc about two times in diameter compared to the protein complex. A systematic determination of the optimal nanodisc size for any given protein of interest will likely aid in optimizing conditions for structural determination. Significant effort has been put forth by previous groups to engineer nanodiscs with sizes ranging from <10 nm to >100 nm, providing a wide range of options for those seeking membrane bound structures. Additional optimization, such as using circularizable nanodiscs that have a more uniform size distribution^48^, may further improve samples for high-resolution characterization.

One striking aspect of our analysis is the observation that AP2 binds to PIP_2_ nanodiscs at a high predict sub-stoichiometric binding using a single-site binding model. For example, for a binding affinity of 1 μM, a complex made at 1 μM of each component has a predicted complex formation of less than 50%. In contrast, crystallographic samples are often used 10-100x more concentrated than cryo-EM, and low-affinity complexes can therefore be driven towards stoichiometric binding by being well above the K_D_ of the interaction. Based on our results, apparent affinities for lipid binding determined using large, planar lipid surfaces like lipid-coated SPR sensors should not be used as a proxy for the affinity of binding to lipid nanodiscs. Rather, researchers seeking to analyze peripheral membrane complexes by cryo-EM should empirically determine if their sample has sufficient affinity to form stoichiometric complexes at the typical concentrations used for cryo-EM, such as by co-elution via gel filtration chromatography. Additionally, other strategies might be used to enhance binding of weakly-associated proteins, such as poly-histidine tags that will bind tightly to nickel-NTA-conjugated lipids. Further development of modified nanodiscs that tether proteins with moderate membrane-binding affinity is therefore of critical importance to visualize these protein-membrane complexes at high resolution.

## Materials and Methods

### Reagents

Protease inhibitors and affinity resins were purchased from Gold Biotechnology, Inc.. 1,2-dioleoyl-sn-glycero-3-phosphocholine (DOPC), 1,2-dioleoyl-sn-glycero-3-phospho-L-serine (DOPS), and L-α-phosphatidylinositol-4,5-bisphosphate (PIP_2_; Brain; Porcine)) were all purchased from Avanti Polar Lipids. Lipidated YxxΦ-cargo peptide (Oleic acid-S{Lys(FITC))}KVTRRPKASDYQRL) was synthesized by Biomatik. HRV3C protease, TEV protease, and benzonase were purified in house.

### Buffers

TBS: 20 mM Tris pH 7.6, 150 mM NaCl

Protease Inhibitor cocktail (final concentration): 0.25 mM AEBSF, 50 μM Bestatin, 5 μM E-64, 20 μM Leupeptin, 5 μM Pepstatin A.

AP2 lysis buffer: 50 mM Tris pH 8.0, 500 mM NaCl, 10% glycerol, 10 mM MgCl_2_, 1 mM CaCl_2_, 0.05% benzonase, 1mM phenylmethylsulfonyl fluoride (PMSF), and home-made Protease Inhibitor Cocktail. AP2 wash buffer: 50 mM Tris pH 8.0, 1000 mM NaCl, 0.05% benzonase, 10% glycerol

AP2 storage buffer: 40 mM Tris pH 8.0, 500 mM NaCl, 1 mM DTT

MSP lysis buffer: 40 mM Tris pH 8.0, 500 mM NaCl, 1% Triton (w/v)

MSP wash buffer 1: same as MSP lysis buffer without protease inhibitors

MSP wash buffer 2: 40 mM Tris pH 8.0, 500 mM NaCl, 50 mM sodium cholate, 20 mM imidazole

MSP wash buffer 3: 40 mM Tris pH 8.0, 500 mM NaCl, 50 mM imidazole

MSP elution buffer: 40 mM Tris pH 8.0, 500 mM NaCl, 500 mM imidazole

MSP storage buffer: 40 mM Tris pH 8.0, 500 mM NaCl, 1 mM DTT

Nanodisc assembly buffer: 20 mM HEPES pH 8.0, 100 mM KCl, sodium cholate at 2x the total lipid concentration.

Binding buffer: 20 mM HEPES pH 7.4, 100 mM NaCl, 1 mM DTT

### Recombinant protein purification

#### AP2 complex purification

AP2 complexes were purified as described previously^16^. Briefly, mouse AP2 “core” (*α*_1-621_, β2_1-591_, μ2, σ2) was co-expressed in BL21 *E. coli* and purified with a C-terminal GST tag on β2 in AP2 lysis buffer. Extensive washing with AP2 wash buffer was required to remove nucleic acid contamination. AP2 was eluted from the resin in AP2 storage buffer by cleavage with HRV3C protease, run on a Superose 6 10/300 size-exclusion column (Cytivia), concentrated using spin concentrators (Amicon), and stored at -80° C.

#### MSP purification

MSP variants were expressed in BL21 *E. coli* in Terrific Broth supplemented with appropriate antibiotics. 1 L culture was grown to mid-log phase at 37°C and expression was induced by addition of 1 mM IPTG. After 2-3 hours of incubation, cells were harvested and resuspended in MSP lysis buffer. Cells were lysed by sonication, clarified by centrifugation, and incubated with 5 mL of pre-equilibrated Nickel-NTA agarose resin for 1 hr at 4° C. The resin was collected in a gravity flow column and washed with 500 mL of each MSP wash buffer and eluted in 30 mL of MSP elution buffer. The protein was dialyzed into MSP storage buffer overnight at 4°C, concentrated with spin concentrators (Amicon), and stored at -80° C.

### Nanodisc assembly

Lipid mixtures were made by combining chloroform lipid stocks at a ratio of 75 mol% DOPC, 15 mol% DOPS, 10 mol% PIP_2_. For cargo-containing nanodiscs, oleic acid-conjugated YxxΦ-cargo was included at 2 mol% and DOPC was reduced to 73 mol%. Lipid films were made by evaporating the chloroform solvent using nitrogen gas and incubating the lipid film under vacuum for >3 hours. Lipid films were hydrated in nanodisc assembly buffer for 30 minutes at 37°C with gentle vortexing every 5 mins. Hydrated films were bath sonicated in 2 min intervals until the solution became transparent, indicating that all lipid had been detergent solubilized. At this point, detergent-solubilized lipids were mixed with an experimentally determined ratio of MSP protein (MSP1 – 1:40; MSP1E3D1 – 1:140; MSP2N2 – 1:160). The solution was allowed to incubate for 20 minutes at room temperature, followed by addition of 0.5 mg of BioBeads (BioRad) that had been washed in methanol, ethanol, and water. The BioBead-lipid-MSP mixture was incubated in a tube rotator overnight at room temperature. Nanodiscs were purified using a Superose 6 10/300 size-exclusion column (Cytivia) pre-equilibrated with binding buffer. Peak fractions were pooled and concentration determined using the absorbance reading at 280 nm.

### Negative Stain Microscopy

CF-200-Cu grids (Electron Microscopy Services) were hydrophilized using a PELCO easiGlow 91000 glow discharge cleaning system. 3 μL of sample at 100 nM were added to the grids and allowed to sit for 30 seconds. Grids were blotted manually using Whatman paper, followed by staining with 1% (wt/vol) uranyl acetate. Excess uranyl acetate was blotted and grids were allowed to air dry. All images were acquired at 120 keV using a FEI Tecnai T12 Transmission electron microscope equipped with a Gatan Rio 16 CMOS camera. Nanodisc diameters were measured manually using FIJI and plotted/ analyzed in PRISM (GraphPad Software).

### Mass photometry

Imaging coverslips (24 × 50 mm^2^ Thorlabs) were water bath sonicated in a 50% isopropanol 50% Milli-Q-H_2_0 solution for 5 minutes, followed by an H_2_O only sonication step (5 minutes). The coverslips were dried using a stream of filtered air. Adhesive, four-well gaskets were pressed onto the clean coverslips to assemble imaging wells (Thorlabs). Prior to imaging, binding buffer was added to the gaskets to determine the imaging plane using an internal autofocusing system. Nanodiscs were added to the imaging buffer at concentrations between 5-15 nM (final volume 20 μL). Measurements were acquired for 1 minute at a frame rate of 100 fps using AcquireMP software (Refeyn Ltd, Oxford, UK). Every sample was independently measured three times (n =3). All measurements were taken on a Refeyn One Mass Photometer (Refeyn Ltd, Oxford, UK) and analyzed using DiscoverMP software (Refeyn Ltd, Oxford, UK) according to previously established protocols.

### Binding assays

5 μM AP2 was mixed with 5 μM nanodisc in a final volume of 200 μL and incubated at room temperature for at least 1 hour. Final salt concentration in the sample was always 100 mM NaCl. Samples were applied to a Superose 6 10/300 size-exclusion column (Cytivia) pre-equilibrated with binding buffer. Binding was verified by comparing elution profiles of AP2 alone, nanodisc alone, and AP2 + nanodisc samples.

### Cryo-EM sample preparation

5 μM AP2 was mixed with 5 μM nanodisc in a final volume of 200 μL and incubated at room temperature for at least 1 hour. The sample was fractionated using a Superose 6 10/300 size-exclusion column (Cytivia) and peak fractions containing AP2 and MSP, as verified by SDS-PAGE, were pooled and concentrated. Samples were applied at 1-2 μM on Quantifoil R 1.2/1.3 300 mesh or Quantifoil R 0.6/1.0 300 mesh grids that had been hydrophilized using a Tergeo-EM Plasma cleaner (PIE Scientific) with a defined Ar/O_2_ gas mixture. Samples were vitrified using a Vitrobot Mark IV (FEI) plunge-freezing device operating at 4° C, 100% humidity and a liquid:propane mixture held at -190 C with liquid nitrogen. Double applications of 3 μL sample typically yielded the highest number of particles in the ice.

### Cryo-EM data collection and processing

All data were collected on a Talos Arctica 200 keV cryoTEM (ThermoFisher) equipped with a K3 direct electron detector (Gatan). Data collection settings were as follows: magnification was either 36,000x (1.11 Å/pixel) or 45,000x (0.88 Å/pixel); defocus was set between -0.6 μm and -1.5 μm; electron fluence rate was 13-15 e^-^/Å^2^/s; total fluence was 45-55 e^-^/Å^2^; 60 total frames. The microscope was operated with SerialEM^49,50^ and the data collection scheme used beam-image shift with a 5×5 collection pattern according to^51^. Beam tilt compensation was calibrated in SerialEM to reduce the residual phase error from large beam-image shifts. Processing was performed using cryoSPARCv3^52^ and Relion4-beta^53^.

Dose-fractionated movies were aligned in cryoSPARC using dose-weighting and patch alignment. Defocus values were estimated in cryoSPARC using the patch function, and micrographs with CTF fits worse than 6 Å and with defocus values outside of 0.5 μm - 2.5 μm were discarded. Particles were picked in crYOLO^54^ using the general model gmodel_phosnet_202005_N63_c17.h5 and in cryoSPARC using the Blob Picker and Template Picker tools. Particles sets were extensively classified using 2D classification, merged, and duplicate particles removed before *ab initio* model generation. 3D classes showing characteristic AP2 density were further 3D classified iteratively between cryoSPARC and Relion before final refinement in cryoSPARC using Non-Uniform Refinement. Maps were *B*-factor sharpened in cryoSPARC and deepEMhancer^55^. Differences in processing for the six total data sets are noted in Supplementary Figs 3-6.

For the MSP2N2 + YxxΦ-cargo dataset, the initial consensus refinement achieved a resolution of 3.3 Å using Non-Uniform refinement in cryoSPARC, although the best regions of the map were those furthest from the membrane binding motifs. To increase the resolution of these regions of the map, focused refinement was performed in cryoSPARC using the Local Refinement tool using three masks generated by manually segmenting the initial refinement using UCSF Chimera^56^. This significantly improved the density for the *α*-σ2 region, allowing for visualization of a bound PIP_2_ headgroup, and the μ2-CTD, allowing for visualization of the bound YxxΦ-cargo peptide. A composite map was generated in Phenix^57^ using the phenix.combine_focused_maps function. Additionally, residue C99 on the σ2 subunit appears to be modified or potentially bound to a cofactor. Oxidation of cysteine residues has been reported as a byproduct of traditional blotting and plunge-freezing methods for cryo-EM grid prep^58^, although this is mitigated by the inclusion of DTT, which is included in our samples. Micrograph number, initial and final particle counts, and refinement statistics for all six datasets are included in Table 1.

Raw data are deposited in EMPIAR under accession numbers EMPIAR-XXXXX, EMPIAR-XXXXX, EMPIAR-XXXXX, EMPIAR-XXXXX, EMPIAR-XXXXX, and EMPIAR-XXXXX.

Maps are deposited in the EMDB under accession numbers EMD-XXXXX, EMD-XXXXX, EMD-XXXXX, EMD-XXXXX, EMD-XXXXX, and EMD-XXXXX.

### Model building and validation

The PDB model of AP2 bound to the YxxΦ motif of Tgn38 (2XA7.pdb^11^) was docked into the MSP2N2 + YxxΦ-cargo AP2 composite structure. The μ2-CTD of this model has a MYC-tag in one of its loops and was therefore replaced with 1BXX^25^, the X-ray crystal structure of the isolated μ2-CTD bound to the YxxΦ motif of Tgn38. Individual subunits were further docked using the “Fit-in-Map” feature of UCSF Chimera. After one round of real-space refinement in Phenix, the model was visually inspected in Coot^59^ and regions of significant deviation from the density were manually rebuilt. At this point, a PIP_2_ ligand was manually placed into the map and another round of phenix.real_space_refine was performed, with secondary structure restraints enabled as well as distance restraints between phosphate groups and AP2 residues to prevent formation of inappropriate linkages. Models were validated by MolProbity^60^ and EMRinger^61^ in Phenix. The final molecular model is deposited in the PDB under accession code XXXX.pdb.

## Acknowledgments

We acknowledge Jared Peck and Dr. Joshua Strauss of the UNC CryoEM Core Facility for technical assistance in this project. We acknowledge Dr. Nathan Nicely of the UNC X-ray crystallography core for technical assistance with mass photometry in this project. We thank Dr. Amy Gladfelter from UNC Chapel Hill for the use of reagents and equipment. We thank Drs. Gunther Hollopeter, Morgan DeSantis, and Jon Fay for thoughtful suggestions to improve the manuscript.

## Author contributions

Conceptualization: KSC, RDS, RWB

Methodology: KSC, RDS, RWB

Investigation: KSC, RDS, KZ, RWB

Visualization: KSC, RDS, RWB

Project administration: RWB

Supervision: RWB

Writing – original draft: RWB

Writing – review & editing: KSC, RDS, RWB

