## Supplemental Figures for "Lipid nanodiscs as a template for high-resolution cryo-EM structures of peripheral membrane proteins"

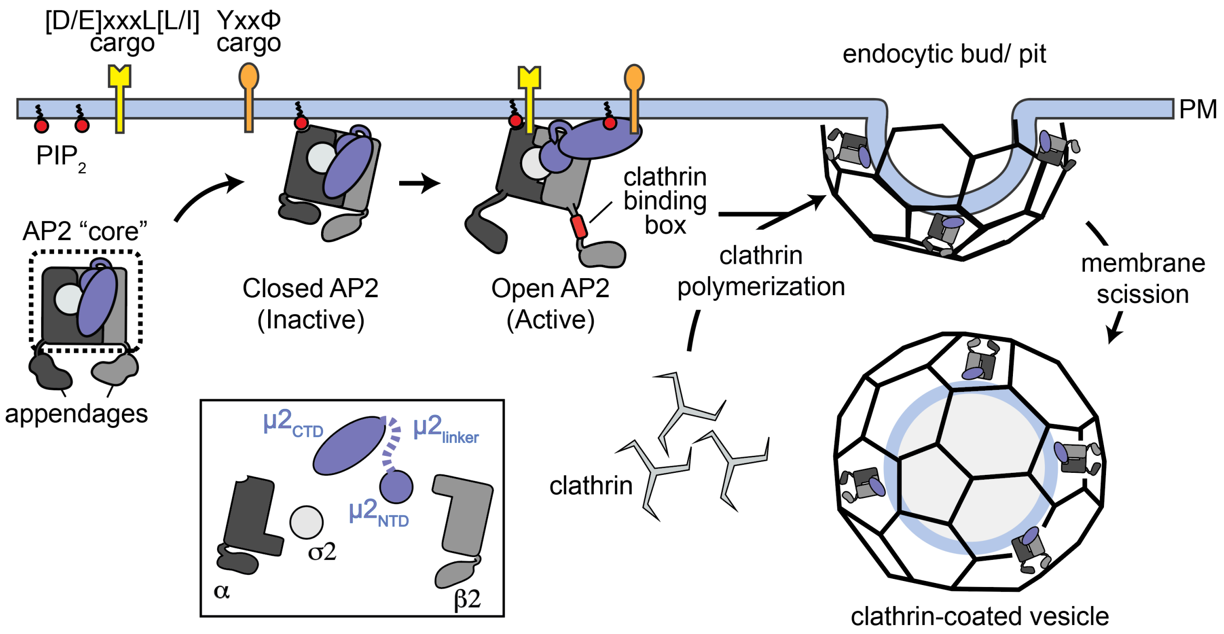


**Supplemental figure 1. Schematic for formation of an AP2-clathrin coated vesicle.** AP2 is a heterotetrameric complex that is largely required for formation of clathrin coated vesicles (CCVs) at the plasma membrane (Inset: schematic of AP2 subunits). AP2's two largest subunits also have "appendages" that are connected to the "core" via unstructured linkers. Most studies, including this one, use AP2 "core" purified recombinantly from *E. coli.* AP2 is recruited to the plasma membrane through intrinsic affinity for $PIP_2$ lipids, which are enriched in this organelle. At the membrane, AP2 undergoes a conformational change, exposing additional $PIP_2$ binding sites and two binding sites for trans-membrane cargo. AP2 recognizes both acidic dileucine ([DE]xxxL[L/I]) and tyrosine-based (YxxΦ) sorting signals, where x represents any amino acids and Φ represents a bulky hydrophobic residue. After engaging cargo, clathrin is recruited via interaction with the AP2 appendages and polymerization leads to membrane deformation. In conjunction with dynamin (not pictured), a coated vesicle is released.

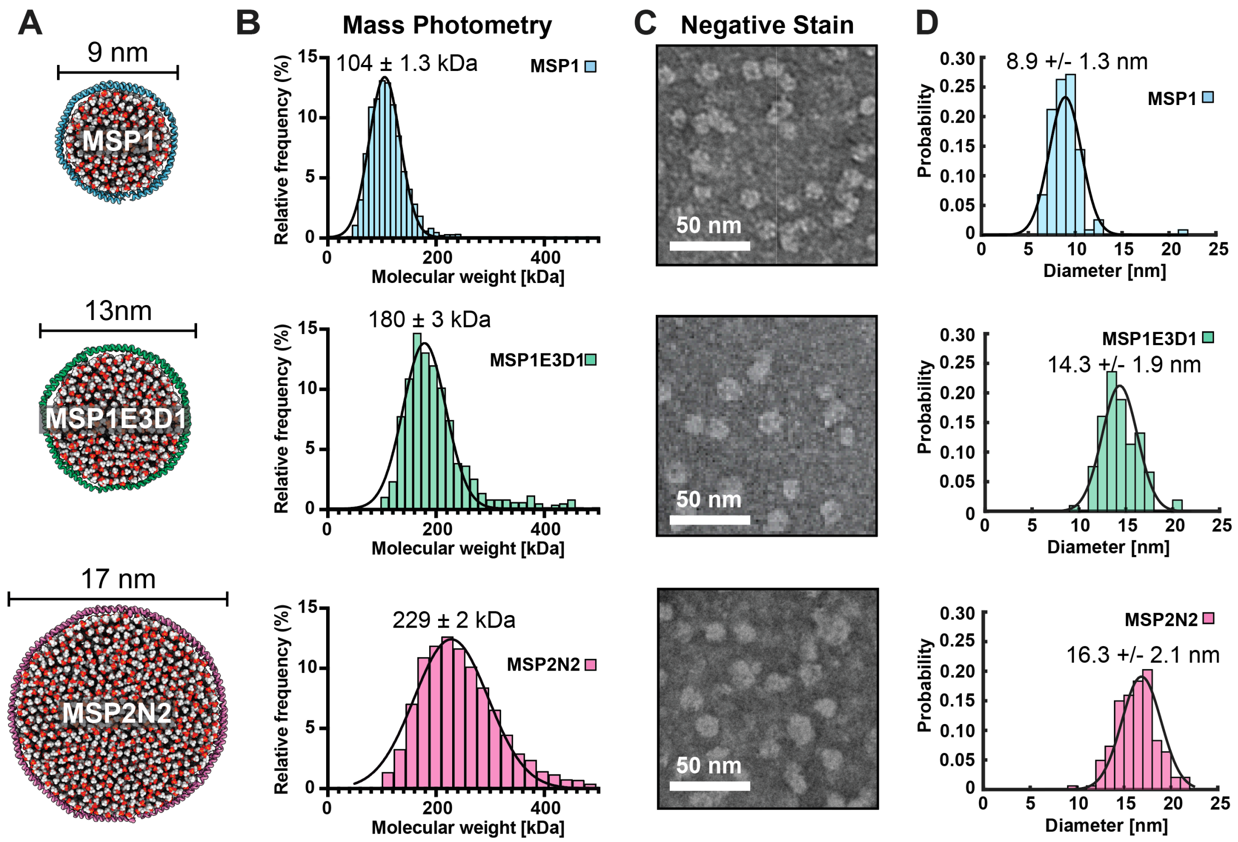


**Supplemental figure 2. Biophysical characterization of nanodiscs.** (A) Molecular models of the three types of nanodiscs used in this study. Models were made to scale. (B) Mass photometry profiles MSP1 (blue), MSPE3D1 (green), and MSP2N2 (pink) nanodiscs. Guassian fitting was used to calculate the mean. N > 4000 counts. (C) Representative negative stain transmission electron microscopy images of each nanodisc. (D) Probability distribution of nanodisc diameters measured from negative stain images. Gaussian fitting was used to calculate the mean. N > 100 nanodiscs per condition.

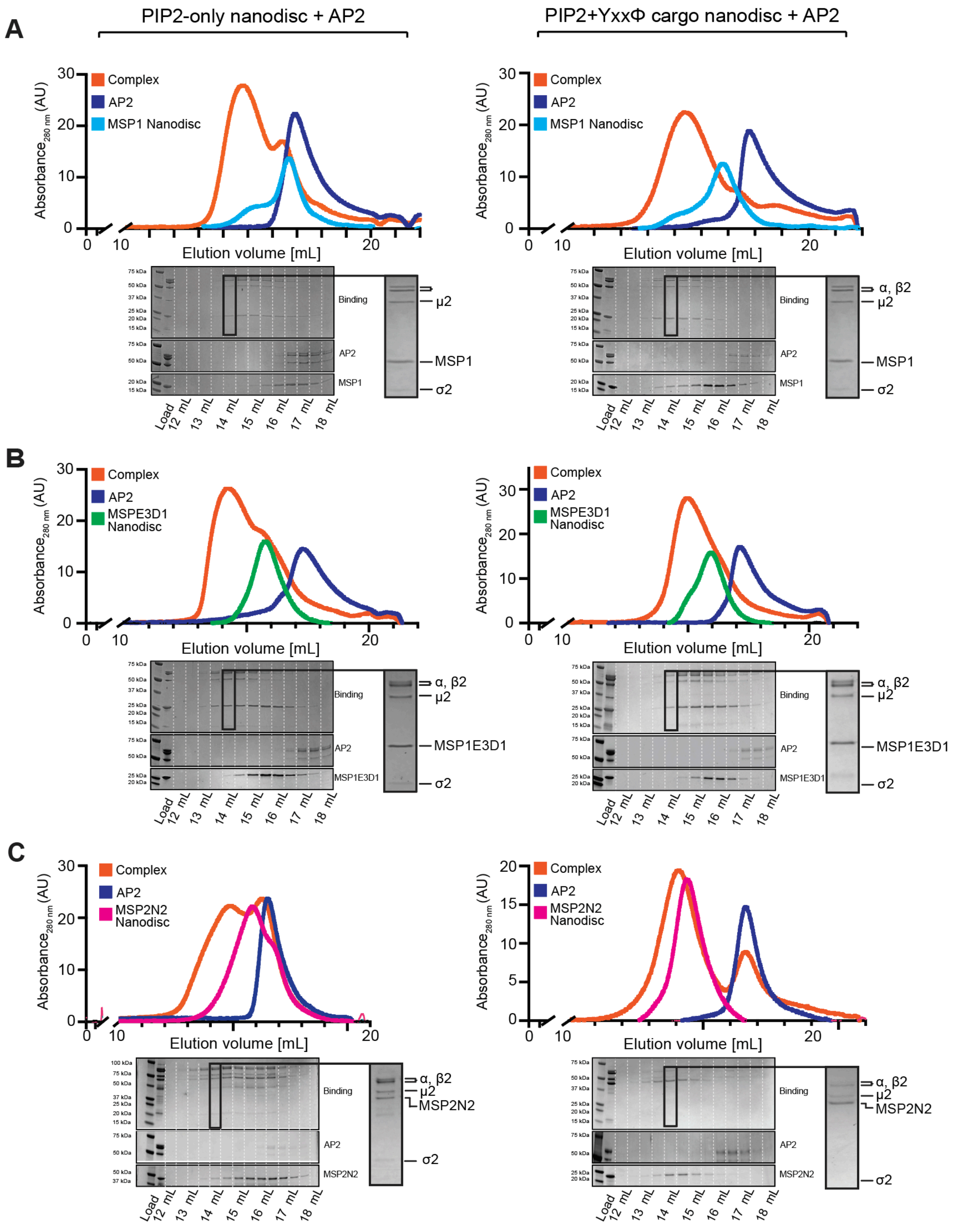


**Supplemental figure 3. Purification of AP2-nanodisc complexes using size exclusion chromatography.** (A-C). Elution profiles of AP2 nanodisc complexes (orange lines) , AP2 alone (purple lines), and nanodisc alone (cyan, green, and pink lines for MSP1, MSPE3D1, and MSP2N2, respectively). Left panels are AP2 binding to nanodiscs without tyrosine cargo embedded in the membrane. Right panels are experiments performed with tyrosine cargo embedded within the membrane. Below each elution profile are the corresponding SDS-PAGE gels.

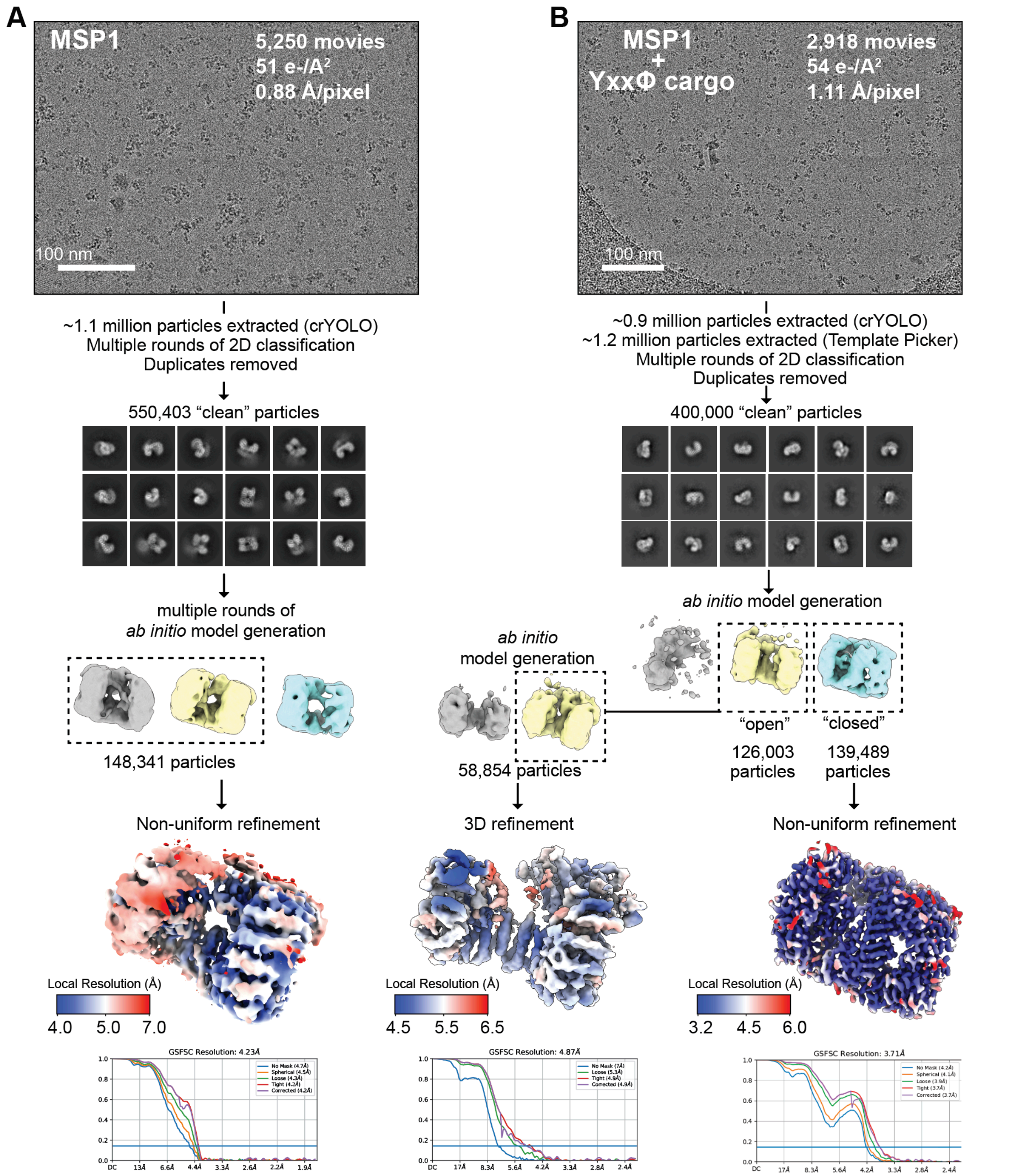


**Supplemental figure S4. Data collection and processing of AP2-MSP1 nanodisc complexes.** A-B) Representative cryo-EM micrographs and processing of AP2-MSP1 nanodiscs with and without YxxΦ-cargo peptide (left and right panels, respectively). Particles were picked using crYOLO and cryoSPARC, followed by several rounds of 2D classification. Selected 2D class averages were used for *ab initio* model generation. Selected *ab initio* models were used for 3D refinement to obtain final cryo-EM maps of the AP2-nanodisc complexes. Shown below each cryo-EM map are standard FSC plots.

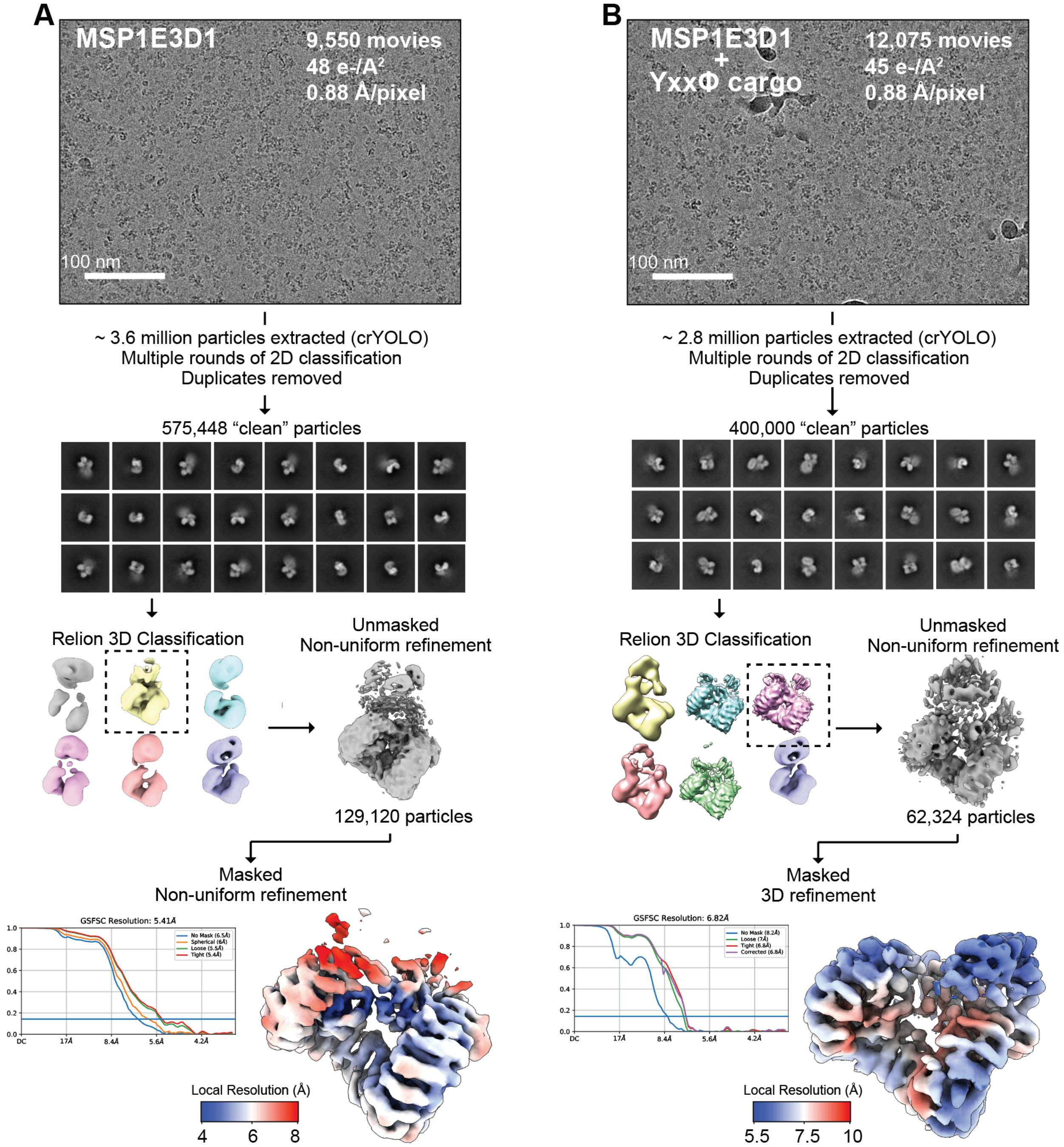


**Supplemental figure S5. Data collection and processing of AP2-MSPE3D1 nanodisc complexes.** Representative cryo-EM micrographs and downstream processing of AP2-MSPE3D1 nanodiscs with and without YxxΦ-cargo peptide (left and right panels, respectively). Particles were picked using crYOLO and cryoSPARC, followed by several rounds of 2D classification. Selected 2D class averages were used for *ab initio* model generation. Selected *ab initio* models were used for 3D refinement to obtain final cryo-EM maps of the AP2-nanodisc complexes. Shown below each cryo-EM map are standard FSC plots.

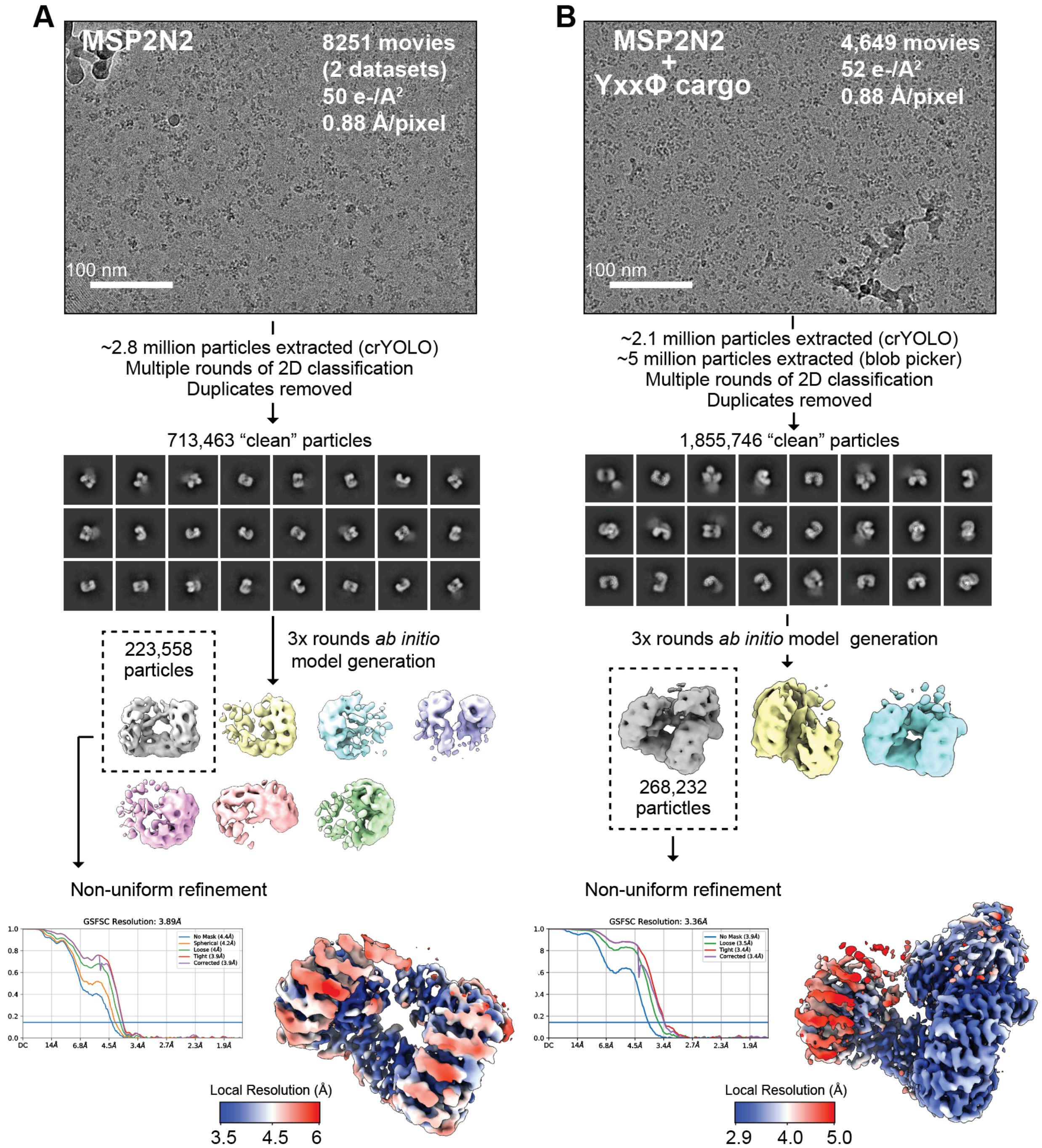


**Supplemental figure S6. Data collection and processing of AP2-MSP2N2 nanodisc complexes.** Representative cryo-EM micrographs and downstream processing of AP2-MSP2N2 nanodiscs with and without YxxΦ-cargo peptide (left and right panels, respectively). Particles were picked using crYOLO and cryoSPARC, followed by several rounds of 2D classification. Selected 2D class averages were used for *ab initio* model generation. Selected *ab initio* models were used for 3D refinement to obtain final cryo-EM maps of the AP2-nanodisc complexes. Shown below each cryo-EM map are standard FSC plots.

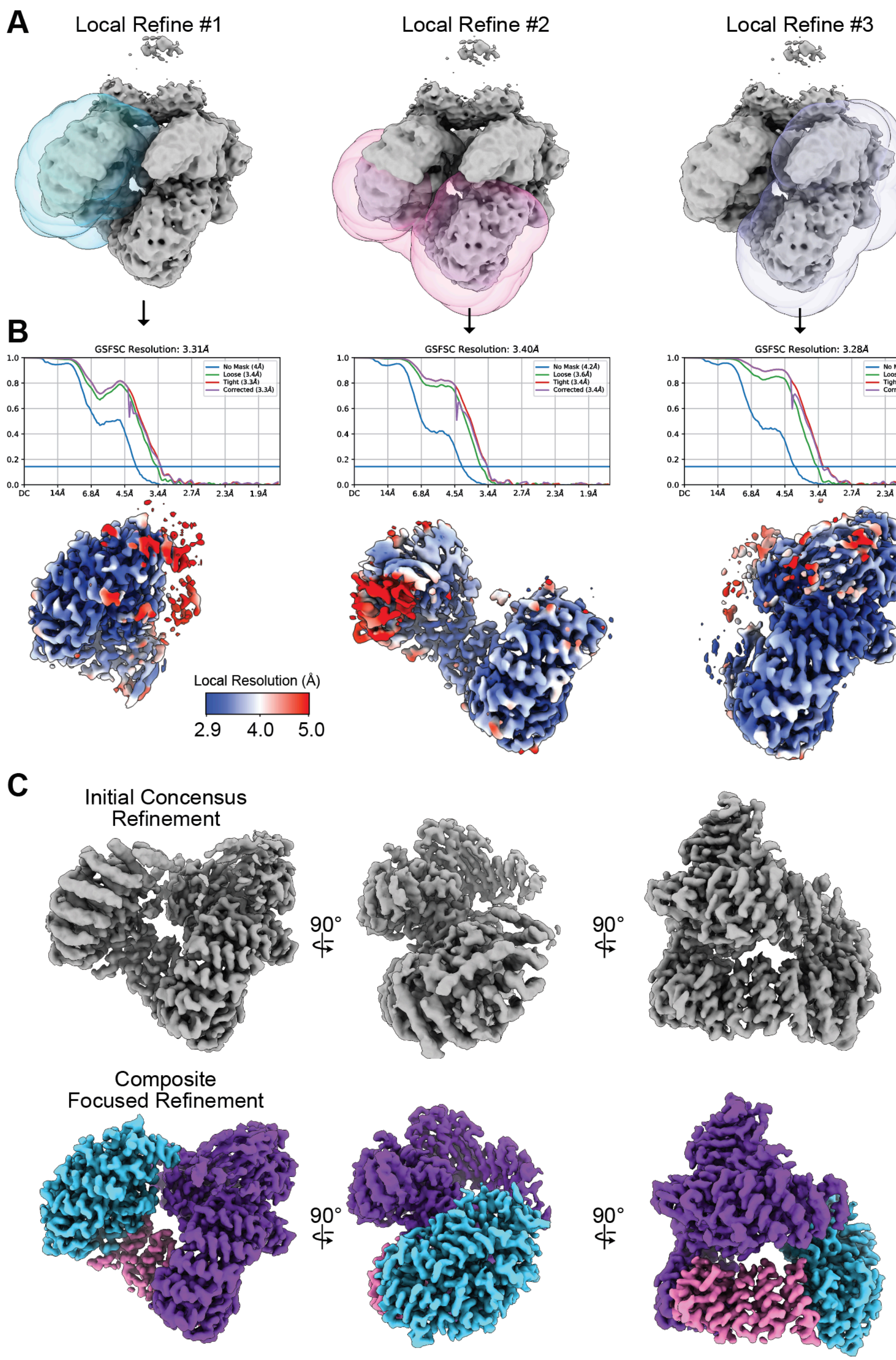


**Supplemental figure S7. Focused refinement of AP2-MSP2N2 + cargo reconstruction.** (A) Local refinement masks for three refinements are shown overlaid on the initial consensus refinement. (B) FSC curves and sharpened 3D volumes colored by local resolution are shown for the three local refinements in (A). (C) Comparison of the consensus refinement (top, grey) and the composite map after local refinement (bottom, colored).

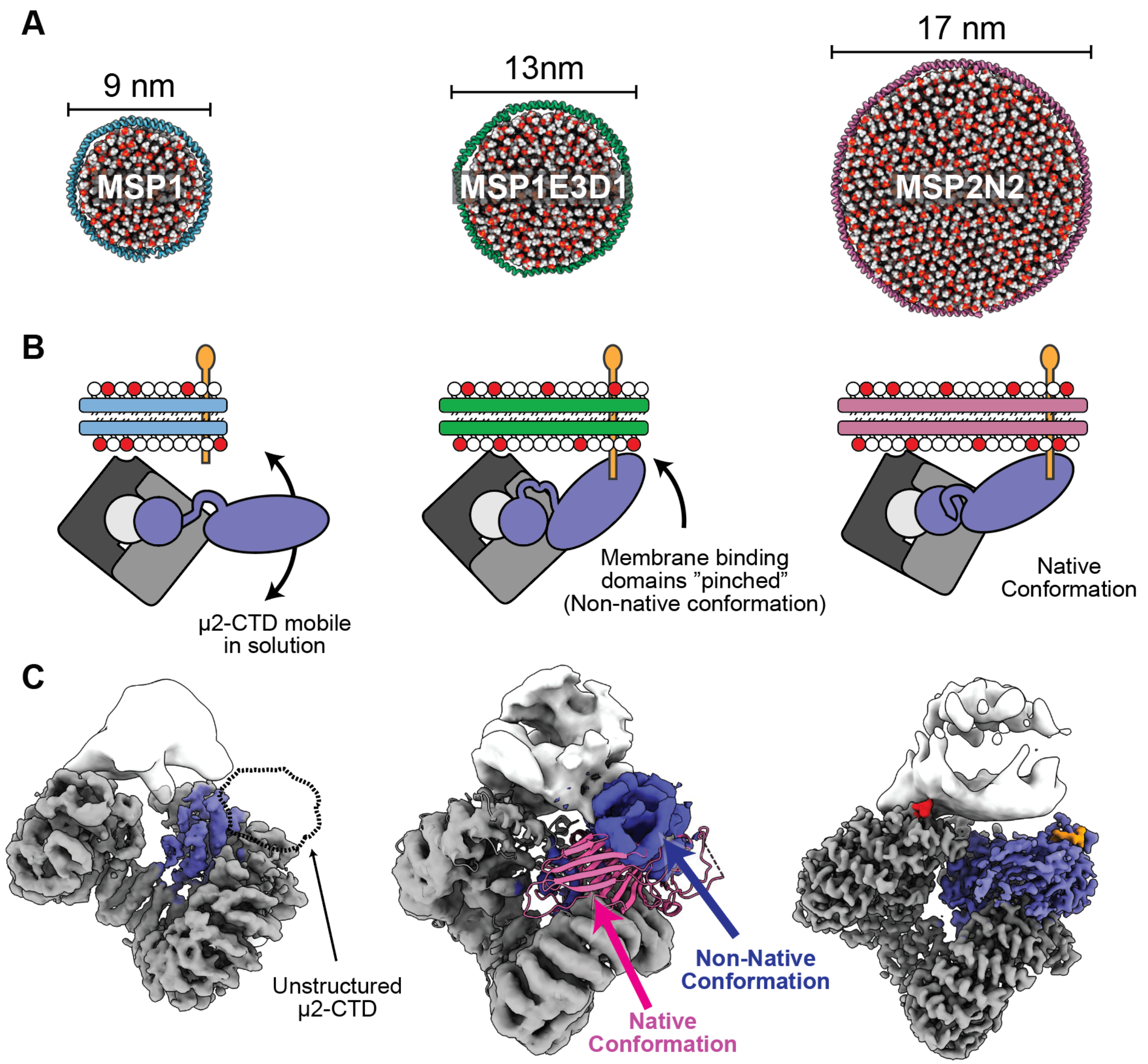


**Supplemental figure S8. Effects of nanodisc diameter on AP2 conformation.** (A) PDB model of the three nanodisc constructs used in this study, with diameter indicated. (2) Cartoon schematic of the three AP2-cargo-nanodisc complexes used in this study. Nanodiscs scaled equally to one another. AP2 not shown to scale. (C) Cryo-EM densities of the three AP2-cargo-nanodisc complexes determined in this study. The missing density for the µ2-CTD in the MSP1 structure is highlighted with a dashed outline. The crystal structure of AP2 + YxxΦ-cargo (2XA7.pdb) is shown docked in the AP2-MSP1E3D1 + YxxΦ-cargo cryo-EM structure. The PDB model is colored pink, showing that the µ2-CTD in the cryo-EM structure on an MSP1E3D1 nanodisc is in a different conformation from the native conformation shown in the crystal structure.
